# Martini 3 Protein Models - A Practical Introduction to Different Structure Bias Models and their Comparison

**DOI:** 10.1101/2025.03.17.643608

**Authors:** Thilo Duve, Liguo Wang, Luís Borges-Araújo, Siewert J. Marrink, Paulo C. T. Souza, Sebastian Thallmair

## Abstract

Biophysical characterization of protein structure and dynamics is essential in many scientific fields, including molecular biology, drug discovery, and enzyme design. Molecular dynamics (MD) simulations have become an increasingly important tool for studying these properties. This chapter provides a hands-on introduction to protein modeling with the Martini 3 coarse-grained (CG) force field. We outline its two-layer framework, where the first layer defines molecular topology and interactions, while the second layer applies structural bias to maintain secondary, tertiary, and quaternary structures. Three structure bias approaches - Elastic Network (EN), GōMartini, and OLIVES - are discussed, highlighting their advantages and trade-offs. Using a protein kinase as a case study, we demonstrate the step-by-step setup of Martini 3 protein models for simulations with the program package GROMACS, including system preparation, fine-tuning, and validation against atomistic reference simulations. Additionally, we explore the combination of an intrinsically disordered region (IDR) with a structured protein domain. By the end of this chapter, readers will have the necessary expertise to apply Martini 3-based protein modeling to a wide range of research applications.

## 1 Introduction

A detailed biophysical characterization of proteins including their structural dynamics is crucial for a broad variety of research fields ranging from molecular biology and drug development to enzyme design and environmentally friendly chemistry. In recent years, modelling techniques became more and more routine tools, offering a complementary approach to experimental techniques for studying protein biophysics. Atomistic and coarse-grained (CG) molecular dynamics (MD) simulations are among the key techniques applied in the field^1,2^. In this chapter, we will provide a hands-on introduction to the protein models available in the fully re-parametrized general-purpose Martini 3 CG force field^3^. We assume that the reader has hands-on experience with the program package GROMACS^4^, for which excellent tutorials are available^5^.

Martini 3 combines top-down and bottom-up coarse-graining with a general mapping of 4-to-1 non-hydrogen atoms to one regular (R) bead^3^. Small (S, 3-to-1 mapping) and tiny (T, 2-to-1 mapping) are also available. The non-bonded interactions are parametrized based on experimental thermodynamic data, while the bonded interactions are parametrized using atomistic^6–8^ or quantum mechanical reference simulations^7,9^. We refer the reader for more details to some recent reviews^10,11^.

The Martini 3 protein model comprises two layers, which are strictly separated from each other: The first layer contains the mapping and chosen chemical bead types as well as the bonded terms; the second layer is the structure bias model. While the first layer corresponds to the standard definition of any molecule in the Martini universe, the second layer is more specific for proteins: To maintain the secondary, tertiary and quaternary structure of proteins, the directionality of interactions, in particular hydrogen bonds, are crucial. Due to the spherical potentials used in Martini 3 and many other CG force fields, this directionality is lost. Therefore, a structure bias model is required to stabilize protein structures. The strict separation between the two layers in Martini 3 enables independent development on both layers and is in contrast to the Martini 2 model^12^, where mappings, bead types, and bonded terms were specific to each of the protein models. The first layer of the current Martini 3 protein model is mostly inherited from the Martini 2 model with moderate updates concerning the new mapping guidelines, bead types, and sizes. The Martini 3 mappings and bead types of the proteinogenic amino acids are depicted in Figure 1. A key difference to the previous model is that the bead type of the backbone (BB) bead does not depend on the secondary structure anymore but is represented by a P2 bead. Exceptions are the special cases glycine (SP1), alanine (SP2), valine (SP2), proline (SP2a), and terminal beads (Q5 for charged and P6 for neutral termini)^13^.

**Figure 1:**
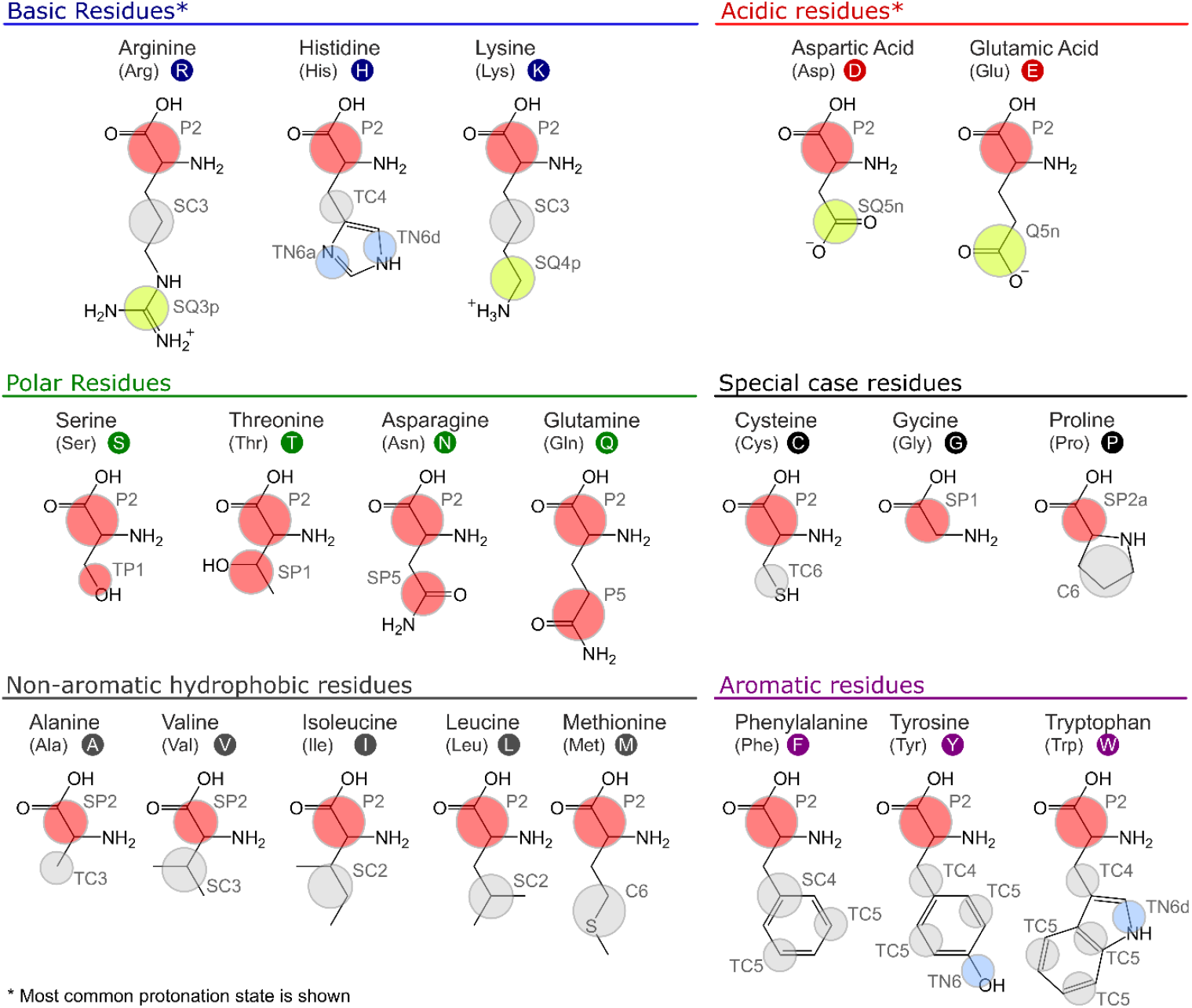
Mapping and chemical bead types of amino acids in the Martini 3 protein model. The main classes of chemical bead types are indicated by colour: P (polar, in red), N (intermediately polar, in blue), C (nonpolar, in grey), and Q (charged, in lime). Different bead sizes are indicated by the radius of the bead, with regular (no symbol), small (S), and tiny (T) beads.

Here, we will focus on the setup, fine-tuning options, and evaluation of the second layer of the Martini 3 protein model – the structure bias model – as well as on dedicated options for intrinsically disordered regions (IDRs). Overall, the Martini protein models require specific bonded terms to model the secondary structure^10,12^, which is in most cases combined with side-chain dihedral corrections^14^. In addition, there are three main options for structural bias models in layer 2 available, typically used to stabilize tertiary and quaternary structures, namely Elastic Network (EN)^3,15,16^, GōMartini^13^, and OLIVES^17^ (Figure 2). Whereas the ad-hoc EN approach provides the most robust and straightforward way of stabilizing a protein structure, and has traditionally been used in Martini, the bioinformatics-based GōMartini approach is nowadays the method of choice as it has proven to be a more versatile method, striking a balance between protein stability and flexibility. Most recently, OLIVES was introduced as a physics-based variant of GōMartini, with the particular prospect of biasing quaternary structures as well.

**Figure 2:**
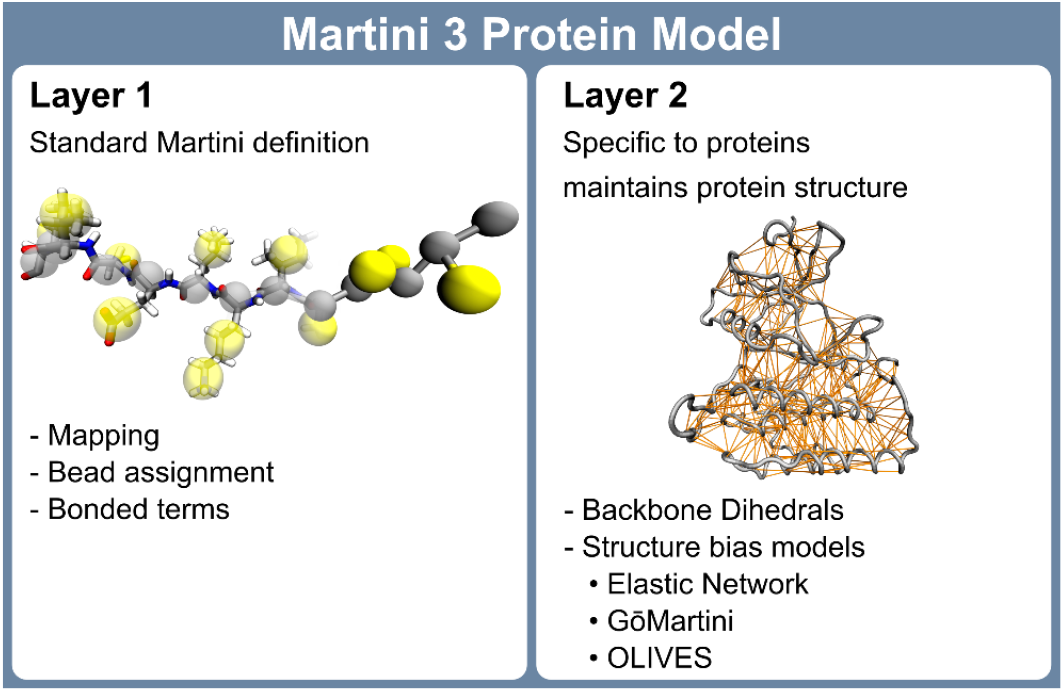
Schematic visualization of the two-layered Martini 3 protein model.

In their simplest form, each of these three structure bias models relies on an atomistic reference, structure which can be obtained from experiments or prediction software. The EN model uses harmonic potentials between the BB beads within a cutoff distance to maintain the protein structure. In contrast, both the GōMartini and OLIVES models employ Lennard-Jones potentials in a Gō-type manner, allowing for contact dissociation. In the GōMartini model, the potentials are defined from a contact map based on native contacts, evaluated through overlap and restricted contacts of structural units^18,19^, combined with a distance cutoff. The OLIVES model, on the other hand, defines its bias by evaluating the hydrogen bonding interactions between BB as well as side chain (SC) beads^17^. For small structured systems, such as single α-helices, the waiver of a structural bias model can be an option as well.

The chapter is structured as follows: Section 2 provides details on how to set up a Martini 3 protein model, taking a protein kinase as our workhorse, and using either EN, GōMartini, or OLIVES, together with secondary structure-specific bonded terms and side-chain corrections. The evaluation of the performance of the model in comparison to atomistic reference simulations is discussed in Section 3, followed by an example for the addition of an IDR to the structured domain of the kinase (Section 4). Additional notes and limitations are discussed in Section 5, before we conclude in Section 6. Related Martini-based tutorial chapters are available for constructing PEGylated proteins^20^, embedding membrane proteins^21^, and for protein-ligand binding^7^.

## 2 Tutorial I - Setup of Martini 3 Protein Structure Models

### 2.1 Preparation of the Protein Structure

In this tutorial, we will prepare different Martini 3 structure bias models for the protein kinase casein kinase delta 1 (CK1δ)^22^. In Sections 2 and 3, we will focus solely on the structured region of this protein (residues 1-292), while the IDR (residues 293-415) will be covered in Section 4. We use an experimental crystal structure as a starting structure^22^. Start by downloading the structure with the PDB code 4JJR from the RCSB-PDB using:

~~~
wget https://files.rcsb.org/download/4JJR.pdb
~~~

To use the PDB structure we first need to remove all atoms that are not part of the protein. This can be water molecules or detergents, ligands, or other molecules used to aid in the crystallization of the protein. The file can be cleaned using the following command.

~~~
grep “^ATOM” 4JJR.pdb > 4JJR_clean.pdb
~~~

Looking at the PDB file, we can see that the protein is present as a dimer. We are interested in the monomer, as CK1δ is a monomer in solution and the dimer is not known to have biological significance^22^. Therefore, we remove chain B with the following command.

~~~
grep “A “4JJR_clean.pdb > 4JJR_clean_A.pdb
~~~

Chain A of the CK1δ structure is missing several residues (43-46, 171-173, 217-222), which need to be modelled before conducting MD simulations. This can be done by providing the SWISS-MODEL web-tool (https://swissmodel.expasy.org/)^23^ with the clean crystal structure (*4JJR_clean_A*.*pdb*) and the respective amino acid sequence. The web-tool returns a PDB structure of the full protein which contains the missing residues. An alternative to using an experimental structure is to predict the full structure of CK1δ using AlphaFold (https://alphafoldserver.com/)^24^. Here, only the amino acid sequence of the protein is needed. Our final structure covers the structured region of residues 1-292.

As a last step in the preparation of our atomistic structure, we check the protonation state of the protein using the H++ web-server (http://newbiophysics.cs.vt.edu/H++/index.php)^25^. Alternatively you can use PROPKA3 (https://github.com/jensengroup/propka-3.0)^26^. We follow the predicted protonation state and use the provided PDB file in the following, which we will name *CK1d*.*pdb*.

### 2.2 Atomistic Reference Simulation

We recommend running atomistic reference simulations to validate the CG Martini 3 protein model (see Section 3). For example, you can use the CHARMM-GUI Solution Builder (https://charmm-gui.org/?doc=input/solution)^27,28^ to build the system and generate topologies using the final PDB file generated in Section 2.1. Solvate the system using 0.15M of NaCl and neutralize it by adding the appropriate number of counter-ions. Here, we recommend conducting at least two replicas of 1 μs each.

### 2.3 Generating the Martini 3 Protein Model

Now that we have a complete atomistic starting structure of CK1δ, we can prepare the CG structure. Here, we use the Martinize2 program (version 0.12.0, https://github.com/marrink-lab/vermouth-martinize)^15^, which generates a CG structure and the necessary topologies to perform simulations with GROMACS based on an atomistic reference structure. The command *martinize2 -h* or the Vermouth documentation (https://vermouth-martinize.readthedocs.io/en/latest/) provide additional information and a complete list of options, which are not discussed comprehensively here. In general, Martinize2 assigns placeholder names to molecules and topology files.

#### 2.3.1 Unbiased Coarse-Grained Model

First, we generate a model without a structure bias model, using the recommended default options for the Martini 3 protein model^13^. This step is done here for illustration purposes only and is normally not recommended for folded domains. We provide Martinize2 with our complete atomistic reference structure using the flag *-f CK1d*.*pdb*. The flag *-ff martini3001* specifies the force field version to be used, in this case, Martini 3.0.0. The *-p backbone* flag generates position restraints of the protein backbone, which are used during the minimization of the system. In addition, the flag *-dssp* adds secondary structure-dependent bonded potentials, designed to stabilize the secondary structure of the protein^29^. By default, Martinize2 uses the mdtraj module^30^ to determine the secondary structure. Alternatively, the DSSP binary can be installed (https://github.com/cmbi/dssp/releases) and provided to Martinize2 via the flag *-dssp path/to/dssp*. Note that Martinize2 is only compatible with DSSP versions 3.1.4 or lower. In older versions of Martinize2, the flags *-cys auto* and *-scfix* need to be specified to add disulfide bridges and to improve the orientations of sidechains^14^, whereas these are the default options as of version 0.12.0.

~~~
martinize2 -f CK1d.pdb -x CK1d_cg.pdb -o CK1d_only.top -ff martini3001 -p backbone
-dssp
~~~

Martinize2 will generate three files: the CG protein structure (CK1d_cg.pdb), the GROMACS topology file (CK1d_only.top), and the protein .itp file (molecule_0.itp).

#### 2.3.2 EN Model

To generate the EN model, the flag *-elastic* needs to be provided, in addition to the previously described flags. The EN model has three parameters, which can be specified using the flags *-ef, -el*, and *-eu* respectively. The flags *-el 0* and *-eu 0*.*85* specify the lower and upper cutoff for the generation of elastic bonds to 0 nm and 0.85 nm, respectively.

~~~
martinize2 -f CK1d.pdb -x CK1d_cg.pdb -o CK1d_only.top -ff martini3001 -p backbone -dssp
-elastic -el 0 -eu 0.85
~~~

Again, three files are created by Martinize2. The EN model added 1004 bonds to the protein topology. To optimize the EN model, the upper cutoff (*-eu*) can be modified. Additionally, the force constant of the harmonic potentials can be changed using the flag *-ef* (see Section 3.1.1). However, we recommend not to use force constants below the default value of 700.0 kJ/(mol nm^2^), as it was shown that lower values cause increased stickiness of proteins^13,31^.

#### 2.3.3 GōMartini

The generation of a Gō-like network using GōMartini necessitates a contact map based on the atomistic input structure^17^, which can be generated using the web-server (http://pomalab.ippt.pan.pl/GoContactMap) with the default settings. Alternatively, the ContactMapGenerator program can be used (https://github.com/Martini-Force-Field-Initiative/GoMartini/tree/main/ContactMapGenerator). In version 0.13.0 of Martinize2, the contact map can also be generated by Martinize2. The contact map is a map of residue overlap (OV) and restricted chemical structural units (rCSU) in the reference structure. The resulting .out file can be downloaded and provided to Martinize2 using the *-go contact_map*.*out* flag. The *-go-moltype* flag enables the user to name the protein and output files. The potential depth of the Gō-like network can be tuned via the *-go-eps* flag. The *-go-low 0*.*3* and *-go-up 1*.*1* flags set the lower and upper cutoffs for contacts which are included in the GōMartini model to 0.3 nm and 1.1 nm, respectively.

~~~
martinize2 -f CK1d.pdb -x CK1d_cg.pdb -o CK1d_only.top -ff martini3001 -p backbone
-dssp -go contact_map.out -go-moltype CK1d -go-low 0.3 -go-up 1.1 -go-eps 8
~~~

The GōMartini model adds 609 potentials to the protein topology. Like the EN model, the GōMartini model can be optimized by modifying the depth of the Lennard-Jones potential (*-go_eps*, see Section 3.1.1).

In addition to the CG protein structure and topology, Martinize2 returns two files: *go_atomtypes*.*itp*, which defines the atom types of the virtual sites, and *go_nbparams*.*itp*, which defines the Gō-like network. It is important **not to change the names of these files**, as they are referenced in other files. We now need to add two *#include* statements to the default *martini_v3*.*0*.*0*.*itp* file. First we include *go_atomtypes*.*itp* in the *[atomtypes]* section using the following command.

~~~
sed -i “s/\[nonbond_params \]/\#ifdef GO_VIRT\n\#include \”go_atomtypes.itp\”\n
\#endif\n\n\[nonbond_params \]/” martini_v3.0.0.itp
~~~

Then *go_nbparams*.*itp* is added to the topology in the *[nonbond_params]* section.

~~~
echo -e “\n#ifdef GO_VIRT \n#include \”go_nbparams.itp\”\n#endif” >>
martini_v3.0.0.itp
~~~

Note, that these commands should only be executed once. It is important that the files go_atomtypes.itp, *go_nbparams*.*itp* and martini_v3.0.0.itp are located in the same directory. Finally, we need to activate the #include statements, by stating *#define GO_VIRT* at the top of the master topology file.

#### 2.3.4 OLIVES

In contrast to the previously discussed protein structure models, the OLIVES model^12^ is recommended to be used without DSSP restraints, to enable conformational changes and partial unfolding of domains. In studies where this is not to be expected one can consider adding DSSP restraints. In our case, we generate two models, OLIVES and OLIVES + DSSP. To create them, we first need to generate a CG protein structure and topology using Martinize2. For OLIVES + DSSP we can use the files we generated in Section 2.3.1. For OLIVES (without DSSP) we use the same command as in Section 2.3.1, but omit the *-dssp* flag.

The OLIVES model is generated using an additional Python script, which can be obtained from GitHub (https://github.com/Martini-Force-Field-Initiative/OLIVES). The respective CG structure and topology files are provided to the OLIVES script using the flags *-c “CK1d_cg*.*pdb”* and *-i “molecule_0*.*itp”*.

~~~
python3 OLIVES_v2.1.1b_M3.0.0.py -c “CK1d_cg.pdb” -i “molecule_0.itp”
~~~

The OLIVES script modifies the provided protein topology files. For CK1δ, the OLIVES model adds 374 potentials to the protein topology. In contrast to the EN and GōMartini models, optimization of the OLIVES model is currently not as well-explored. Nevertheless, there are options available to tune the cutoff for the generation of the secondary and tertiary network via the flags *–ss_cutoff* and *–ts_cutoff* and to scale the strength of the secondary and tertiary structure bonds using the flags *–ss_scaling* and *–ts_scaling*, respectively.

Figure 3 shows the bond networks created for CK1δ by each structure bias model.

**Figure 3:**
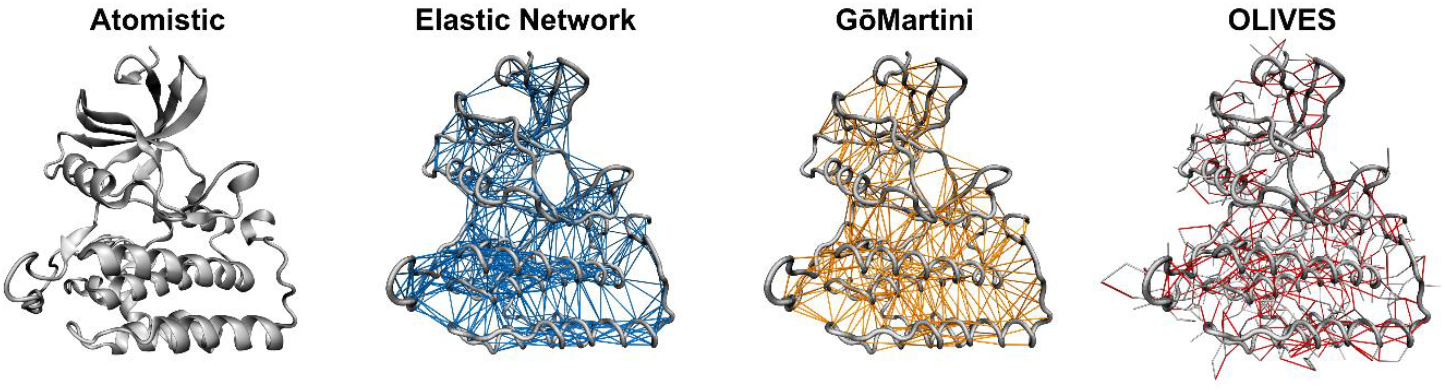
Structure bias models for CK1δ. Atomistic structure in cartoon representation and the bond networks created by EN (blue, 1004 bonds), GōMartini (orange, 609 potentials), and OLIVES (red, 374 potentials) are shown. Note that while the additional bonds in the EN and GōMartini model exclusively connect the backbone beads, also side chain beads are connected in the OLIVES model.

### 2.4 Coarse-Grained Martini 3 Simulations

With the coordinates and topology of the protein, we can prepare a starting system for a simulation by placing the protein in a simulation box. For this task, we will use the Python script insane (https://github.com/Tsjerk/Insane)^32^. Alternatively, one can use *gmx solvate* or COBY (https://github.com/MikkelDA/COBY)^33^. We specify the box type (*-pbc cubic*) and size (*-box 10,10,10*) and solvate the protein in water (*-sol W*), adding 0.15 M of NaCl *(−salt 0*.*15*), while automatically adding the appropriate amount of counter ions to neutralize the system. Note that when using older versions of Martinize2 in combination with the GōMartini model, the charge of the protein needs to be specified using the *-charge 11* flag. As the virtual sites for the GōMartini model were added at the end of the file, the insane script counts each residue twice and therefore doubles the charge.

~~~
insane -f CK1d_cg.pdb -o CG.gro -p system.top -pbc cubic -box 10,10,10 -salt 0.15
-sol W -d 0 -charge 11
~~~

Insane adds NA+ and CL-to the .gro and topology file, a remnant of the Martini 2 nomenclature, while the standard Martini 3 ions are labelled NA and CL. Therefore, remove the charge sign in *CG*.*gro* and *system*.*top*. You may get a list of molecules in the .top file similar to this:

~~~
[molecules]
; name number
CK1d    1
W    8230
NA     85
CL     96
~~~

Additionally, add the relevant Martini 3 force field topologies at the top of the master topology file.

~~~
#define GO_VIRT; only for GoMartini
#include “./martini_v3.0.0.itp”
#include “./martini_v3.0.0_ions_v1.itp”
#include “./martini_v3.0.0_solvents_v1.itp”
#include “./CK1d.itp”
~~~

The system setup is depicted in Figure 4.

**Figure 4:**
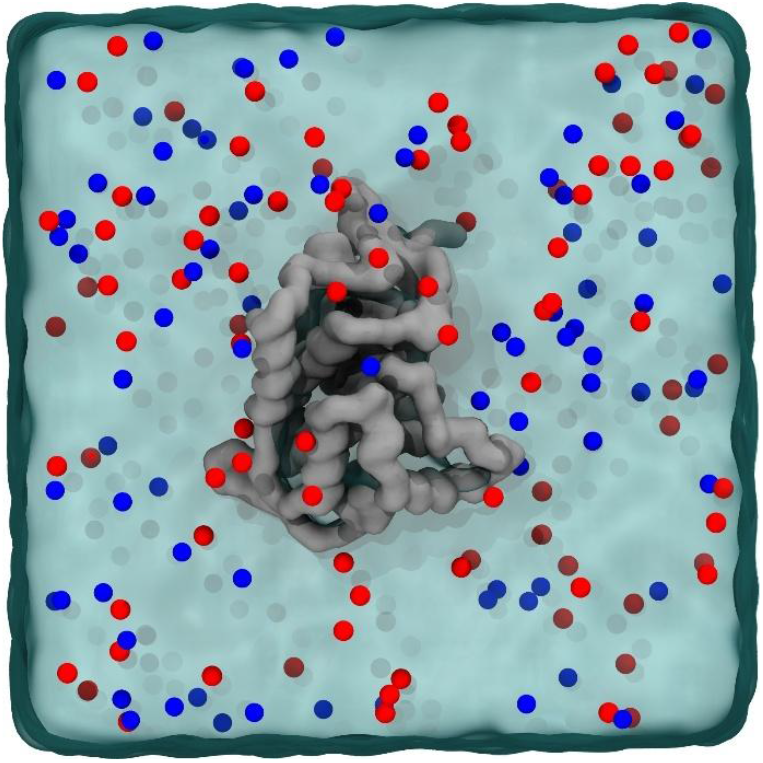
Setup of the simulation box, containing CK1δ (silver), Na+-ions (red) and Cl--ions (blue) solvated in water (blue transparent surface).

We are now ready to perform CG MD simulations of CK1δ using different structure bias models. First, we will do a short energy minimization, followed by a position-restrained NPT equilibration. The previously generated position restraints can be used by specifying *define = -DPOSRES* in the .mdp file.

~~~
gmx grompp -p system.top -c solvated.gro -f minimization.mdp -o minimization.tpr
-r solvated.gro
gmx mdrun -deffnm minimization -v
gmx grompp -p system.top -c minimization.gro -f equilibration.mdp
-o equilibration.tpr -r solvated.gro
gmx mdrun -deffnm equilibration -v
~~~

We can now start a production run. If the simulation crashes, more equilibration steps might be needed. However, it’s important to note that for simple smaller proteins, such as this one, Martini is generally quite robust, and crashes may indicate issues with the built models or systems. Note that you will get a warning about the simultaneous use of the Parrinello-Rahman barostat and newly generated velocities. This can be ignored by using the *-maxwarn 1* flag.

~~~
gmx grompp -p system.top -c equilibration.gro -f dynamic.mdp -o dynamic.tpr
-maxwarn 1
gmx mdrun -deffnm dynamic -v
~~~

Note that when using the OLIVES model, the flags *-noddcheck -rdd 2*.*0* need to be specified in *gmx mdrun* to avoid domain decomposition errors during the simulation^12^. To judge the proficiency of each model to replicate the properties of a given protein, we recommend at least three replicas of 250 ns, following the recommended 1:4 ratio for comparisons between atomistic and CG Martini simulations, due to the smoothed energy landscape of CG simulations ^34^. 250 ns of CG simulations correspond to approximately 1 μs of effective simulation time.

The following section will discuss different metrics that can be used to judge which protein structure model is the best suited for a specific use case.

## 3 Tutorial II - Metrics to Compare Protein Structure Models

### 3.1 Root Mean Square Fluctuation (RMSF)

The Root Mean Square Fluctuation (RMSF) is a metric that quantifies the fluctuation of the position of an atom over time. It is calculated according to the following equation, where ***r***_*i*_ is the coordinate vector of a particle *i*, 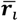 the ensemble average coordinate vector of *i* and T the total number of frames in the trajectory.

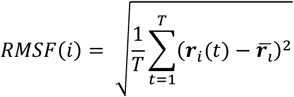

The RMSF can be used to quantify the flexibility of a structure over the course of a simulation. Regions with high RMSF values indicate high structural mobility, while regions with low RMSF values are more rigid.

To compare the RMSF between two simulations, e.g. an atomistic reference and the CG counterpart, the ΔRMSF can be calculated with the following equation. An ideal model would have a ΔRMSF of zero for all residues.

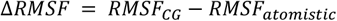

In our case we use a script to calculate the RMSF of the Cα-atoms or backbone beads for atomistic and CG trajectories, respectively. First, the atomistic trajectories are divided into 200 ns segments, while CG trajectories are divided into 50 ns segments, following the 1:4 ratio recommended for comparisons between atomistic and CG Martini simulations, due to the smoothed energy landscape of CG simulations^34^. Subsequently, the protein is centred in the simulation box. In seven iterations, the trajectory is fitted to all C_α_-atoms (atomistic) or BB beads (CG) with a mean RMSF value below the threshold of 1.5 Å in the previous iteration, using *gmx trjconv* with the flag *-fit rot+trans*. This is done to avoid fitting the trajectory to highly flexible regions of the protein. The final mean RMSF values for each residue are calculated following the seventh iteration. RMSF values were computed using *gmx rmsf* with the flags *-nofit -res*. The script used for this calculation can be found at https://cgmartini.nl/.

The RMSF of a protein is a key metric to judge the viability of a protein structure model, as it can indicate to what extent a certain model can reproduce the intrinsic flexibility of a protein. In the following sections, we will first use it to determine the optimal parameters for the EN and GōMartini models. Then, we will compare these optimized models to the OLIVES model.

#### 3.1.1 Tuning of the EN and GōMartini Models

Figure 5 depicts the ΔRMSF for the different parameters of the EN and GōMartini models. We tested three values for the force constant *k*_B_ of the EN model: 700, 800, and 1000 kJ/(mol nm^2^). Overall, only minor changes in the RMSF of the protein can be observed. As expected, a higher force constant decreases the overall flexibility of the protein and therefore the RMSF.

**Figure 5:**
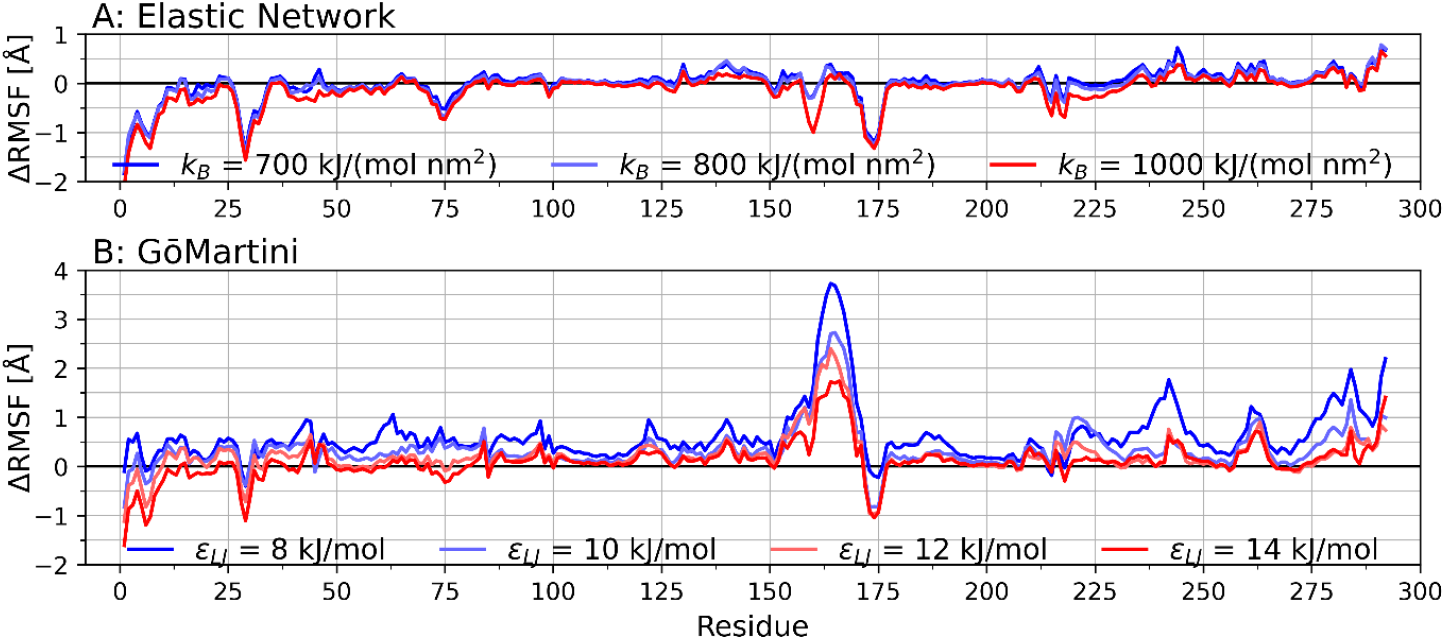
Backbone RMSF deviation (ΔRMSF) between the CG models and the atomistic reference of CK1δ using different parameters for the EN and GōMartini models. A: EN model with kB = 700, 800, and 1000 kJ/(mol nm2). B: GōMartini model with εLJ = 8, 10, 12, and 14 kJ/mol.

Figure 5A shows that the higher force constants, particularly 1000 kJ/(mol nm^2^), lead to an underestimation of the flexibility of CK1δ. Therefore, we will proceed with the force constant value of 700 kJ/(mol nm^2^) in the following steps.

For the GōMartini model, four values for the potential depth *ε*_LJ_ were tested: 8, 10, 12, and 14 kJ/mol. Figure 5B shows that the potential depth of the GōMartini model has a substantial impact on the RMSF of CK1δ. Overall, higher values for *ε*_LJ_ better capture the flexibility of CK1δ, both in more rigid regions, as well as in highly flexible loops. Therefore, we will proceed with a potential depth of 14 kJ/mol.

#### 3.1.2 Comparison of the Structure Bias Models

When comparing the C_α_/BB RMSF of different structure bias models, examine the RMSF on a per-residue basis to identify deviations in specific regions, paying attention to whether the flexibility of the more rigid regions is reproduced. Flexible regions, such as loops and termini, while important, are more challenging to reproduce. Typically, more deviation from the reference is accepted here for the overall evaluation Figure 6 shows the RMSF distribution and RMSF for each residue in comparison to the atomistic reference simulation (as opposed to the ΔRMSF in Figure 5). This allows us to identify highly flexible and more rigid regions of CK1δ. Depending on the scientific question at hand, it might be useful to focus the analysis on certain regions of the protein, which are important for the specific system, e.g. a ligand binding pocket or a protein-protein interface.

**Figure 6:**
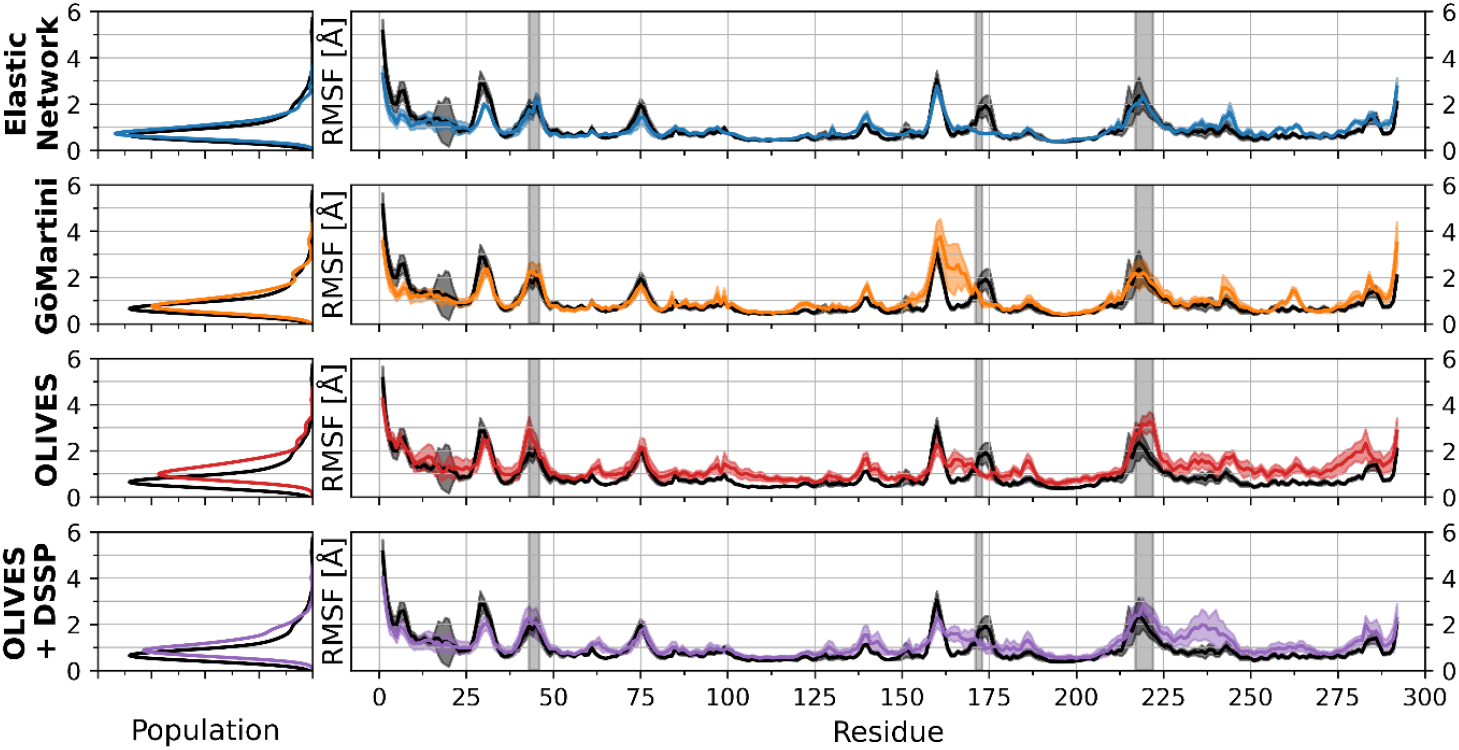
Comparison of the backbone RMSF of CK1δ between protein structure models (EN, GōMartini, OLIVES and OLIVES + DSSP). Left: RMSF distribution over all residues and time windows. Right: RMSF of individual residues. The atomistic reference is displayed in black. Solid lines show the mean of all time windows, the shaded area the mean ± standard deviation. Residues missing in the reference crystal structure are indicated by the shaded rectangles.

With a mean RMSF of 0.91 Å, CK1δ is a rather rigid protein, favouring more rigid protein models, such as the EN model and GōMartini model with a high potential depth (see Section 3.1.2). Figure 6 shows that both the EN and GōMartini model overall capture the flexibility of CK1δ very well, especially in the more rigid regions. However, both models slightly overestimate the RMSF of residues 260-265, while underestimating the flexibility of the loop in the region 171-175. The GōMartini model shows an increased RMSF in the residues 163-171 located between two loops.

In contrast, the OLIVES model consistently overestimates the flexibility of the more rigid regions of CK1δ, but also two of the loops that are missing in the crystal structure (residues 43-46 and 217-222). The addition of DSSP secondary structure restraints resulting in the OLIVES + DSSP model reduces this overestimation and reproduces the flexibility of the rigid region between residues 50-150 and the two loops (residues 43-46 and 217-222) well. However, while improved compared to the OLIVES model, the RMSF of the rigid region between the residues 225-285 is still overestimated with the OLIVES + DSSP model. Like the GōMartini model, both models overestimate the RMSF of residues 163-171. For the OLIVES model, a scaling of the interaction strength is also possible (see Section 2.3.4). This might improve the comparison to the atomistic reference To conclude, in this case both the EN and GōMartini models demonstrate strong performance in reproducing the RMSF of the atomistic reference simulation, with the EN model showing a slight edge. These models especially capture the dynamics of rigid regions most accurately. The combination of the OLIVES model and DSSP restraints provides an acceptable performance, although less precise than the EN and GōMartini models, due to an overall overestimation of the flexibility of CK1δ. The OLIVES model alone struggles to accurately reflect the RMSF of CK1δ, especially in rigid areas.

### 3.2 Root Mean Square Deviation (RMSD)

The Root Mean Square Deviation (RMSD) is a measure used to quantify structural similarity, typically between a frame of a simulation and a reference structure, such as a crystal structure. To calculate the RMSD along a trajectory, at each frame, the structure is aligned to the reference structure. Then, the RMSD is calculated according to the following equation, where ***r***_*i*_(*t*) is the coordinates of a particle *i* at frame t, 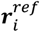 the coordinates of particle *i* in the reference structure, and N the total number of particles.

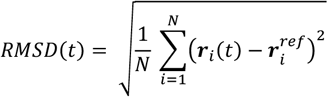

One might notice the similarity between this equation and the equation to calculate the RMSF (see Section 3.1). While both metrics have a similar concept, they contain different information. The RMSD provides a global comparison, assessing overall structural deviations relative to a reference structure over time. In contrast, the RMSF is calculated relative to an averaged structure and measures the flexibility of individual residues over the course of a trajectory. While the RMSD evaluates global structure deviation, the RMSF provides insight into local dynamics and flexibility.

In our case, we calculate the RMSD of all C_α_-atoms (atomistic) or BB beads (CG) along our trajectories. First, we centre the protein in the simulation box. Additionally, we create an index file that contains the indices of all C_α_-atoms (atomistic) or BB beads (CG). Then we use this index file and *gmx rms* to calculate the RMSD. The command will prompt you to first select the index group on which to fit the trajectory, then which index group to use for the calculation of the RMSD. We select CA/BB for both.

~~~
gmx trjconv -f production.xtc -pbc whole -center -s production.tpr -o noPBC.xtc
gmx make_ndx -f equilibration.gro -o index.ndx
gmx rms -f noPBC.xtc -s production.tpr -n CA.ndx -o rmsd.xvg
~~~

It might be useful to also consider other selections for the RMSD calculation. For example, one could calculate the RMSD of a ligand bound to a binding pocket relative to a crystal structure, to judge the ability of a model to reproduce experimentally observed binding modes. The RMSD of the protein can also be used to judge if the system is well equilibrated. For an equilibrated system, one expects a converged RMSD time series.

#### 3.2.1 The Unbiased Coarse-Grained Model

Figure 7 illustrates the need for the structure bias layer in Martini 3. While the atomistic trajectory and biased CG trajectories show RMSD values of below 4.5 Å, the unbiased model exhibits values of up to 23 Å. This corresponds to the unfolding of the protein depicted in Figure 7C, showing that the Martini 3 protein model cannot retain a tertiary structure without additional biasing.

**Figure 7:**
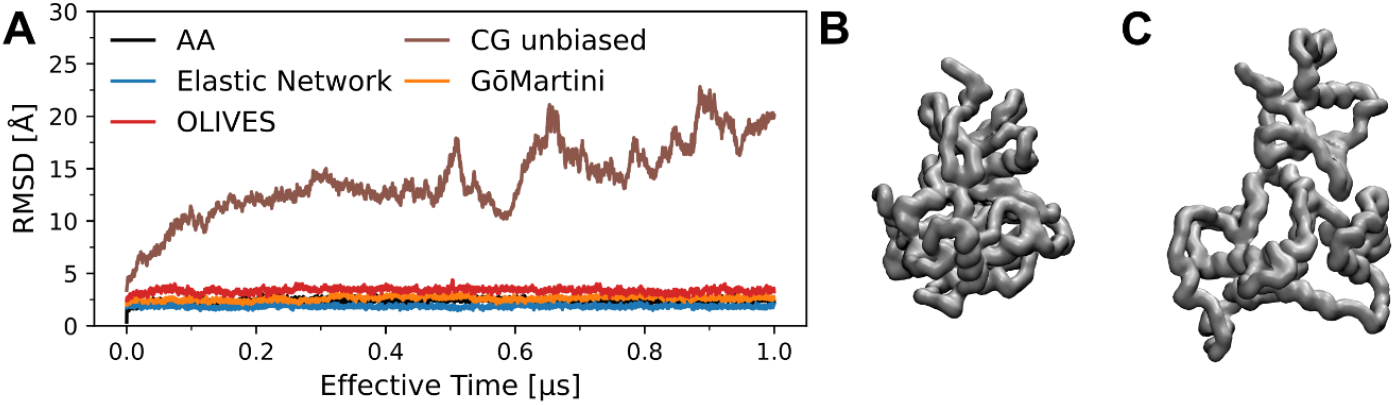
Unbiased CG Model. A: Example RMSD traces over time of atomistic reference and CG protein structure models. B,C: Snapshot of unbiased CG model at the beginning of the simulation (B) and after 0.9 μs effective time (C).

#### 3.2.2 Comparison of the Structure Bias Models

Figure 8 shows the RMSD distributions of the atomistic reference and the respective structure bias models for CK1δ. We see that the atomistic reference distribution ranges from 1.5 – 3.0 Å. The OLIVES model shows the distribution with the highest deviation from the atomistic distribution, ranging from 2.5 – 4.5 Å. In agreement with previous results, the addition of DSSP restraints to the OLIVES model leads to a decrease in the RMSD and higher agreement with the atomistic reference. The GōMartini shows even higher overlap with the atomistic reference, resulting in the best estimation of all models here, while also showing higher RMSD values up to 3.5 Å not present in the atomistic reference. The EN model shows the lowest RMSD values of all models, with values ranging from 1.5 – 2.5 Å. While the distribution overlaps with the atomistic reference distribution, the model is unable to reproduce the higher RMSD values also present in the atomistic reference. This indicates the strong biasing of the EN model towards the crystal structure that was used to create the model.

**Figure 8:**
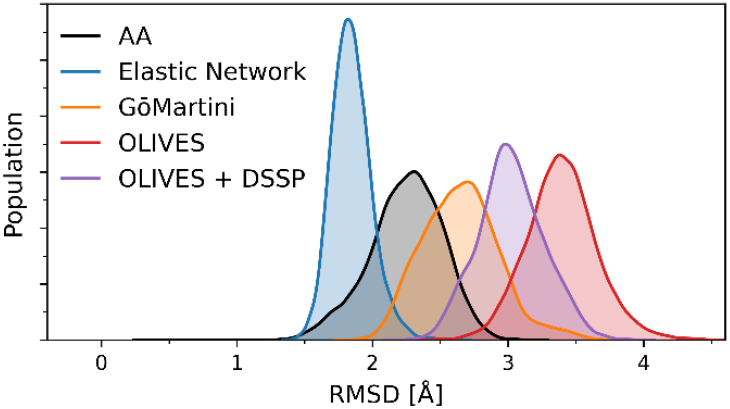
RMSD distributions of CK1δ. Atomistic used Cα-atoms and CG models used BB beads to fit the trajectory and calculate the RMSD.

### 3.3 Backbone Distance Matrix

The final metric we are going to analyse is the distances between the backbone beads of CK1δ. Using a nested loop and *gmx distance* we can calculate this distance for each residue pair.

~~~
gmx distance -f noPBC.xtc -s production.tpr -select “resid ${i} ${j} and name BB”
-len 4.0 -binw 0.01 -oh ${i}-${j}_BB-dist.xvg
~~~

For the reference atomistic trajectory we must ensure that the distance is calculated between where the BB bead would be located, at the center of mass (COM) of the backbone atoms. Using the flag *-select “resid $*{*i*} *$*{*j*} *and name N CA C O”* we select all non-hydrogen atoms of the protein backbone. We then specify that the COM should be used for the calculation using *-selrpos part_res_com* and *-seltype part_res_com*.

~~~
gmx distance -f noPBC.xtc -s production.tpr -select “resid ${i} ${j} and name N CA
C O” -selrpos part_res_com -seltype part_res_com -len 4.0 -binw 0.01
-oh ${i}-${j}_BB-dist.xvg
~~~

This returns a histogram of distances for all residue pairs, ranging from 0 to 80 Å (specified by *-len 4*.*0*). Note that this approach generates a large number of files, so check your storage availability and delete the created files after performing the necessary calculations.

We can now calculate the mean of each histogram and plot them in a matrix. To judge the similarity of the distance distributions of a protein structure model to the atomistic reference, we can calculate the integrated absolute difference^35^. For this, we calculate the absolute difference of two histograms and sum up the values. Identical distributions then have a sum of zero, while distributions with no overlap sum up to two. Figure 9 shows both metrics for all models of CK1δ. Similarly to the analysis of the RMSF, here, the focus should be rather on rigid regions or regions of interest, such as potential ligand binding sites.

**Figure 9:**
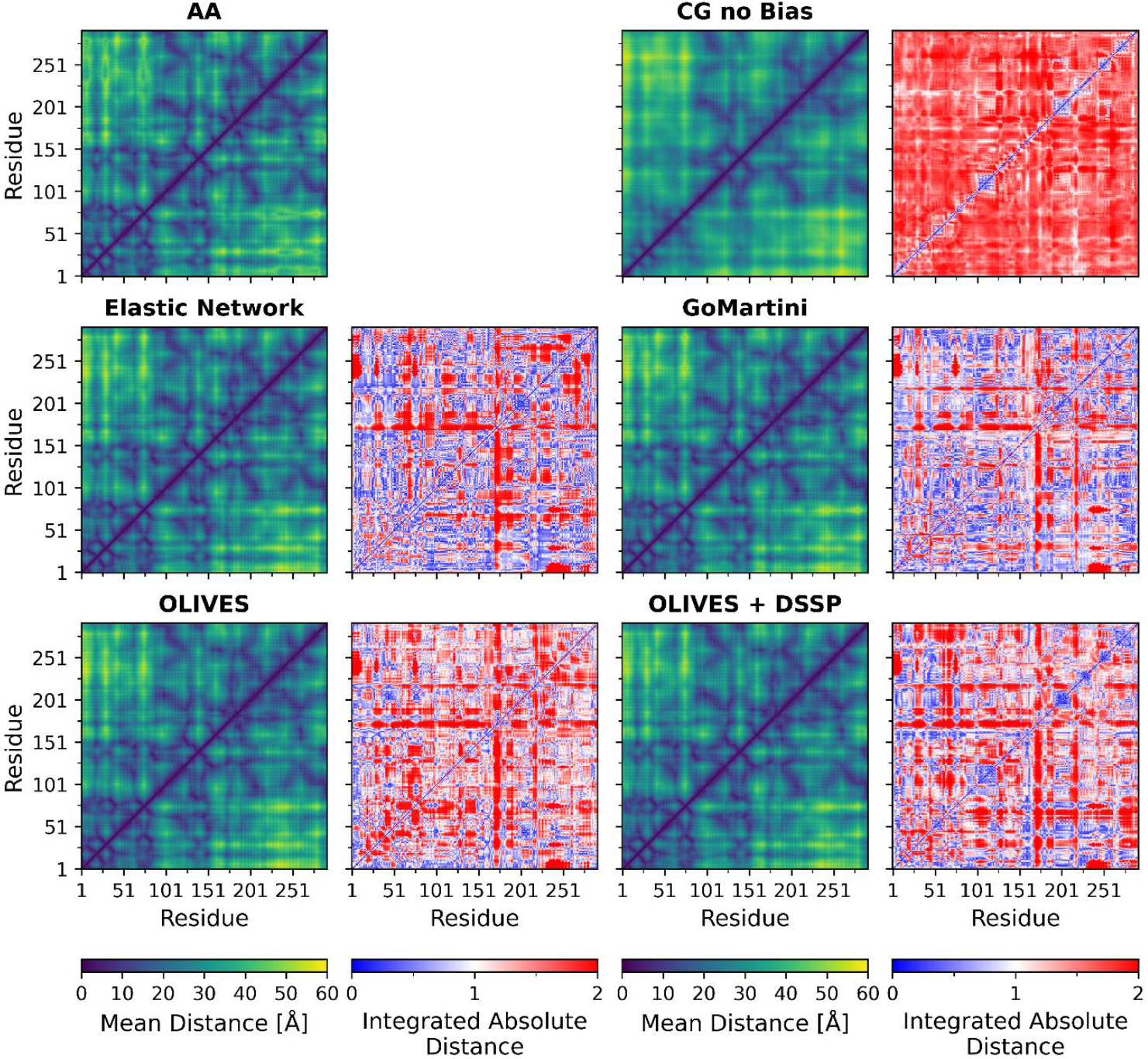
Matrix representation of backbone bead to backbone bead distances for all residues. Displayed is the mean distance and integrated absolute difference.

As we saw with the RMSD in Figure 7, the CG model without a structural bias shows larger distances than the atomistic reference. This is the case for all residue pairs, expect for individual pairs near the diagonal, which represents very short-range interactions mediated by the DSSP restraints. This shows that while the protein structure largely unfolds during the simulation, some secondary structure motifs remain intact.

Notably, in all models the distance pairs including the missing loop in the crystal structure (residues 171-173) show large deviations from the atomistic reference for all models. This is in line with the deviations of the RMSF in this region that was observed for all models (see Section 3.1.2). Interestingly, all models generally displayed deviations between residues 175 and 200, although the EN and GōMartini reproduce the flexibility of this region well. A comparison between the OLIVES and OLIVES + DSSP models reveals that OLIVES + DSSP produces better agreement at distance pairs along the diagonal, corresponding to short-range interactions. This indicates a better representation of the secondary structure of CK1δ. In terms of overall distance accuracy, the GōMartini model outperforms the others, displaying the best overall fit. The highest deviations in distances are within loop regions.

## 4 Tutorial III – Simulations of IDRs in Martini 3

This section focusses on introducing the IDR of CK1δ to the system. Previous works have shown, that simulations of IDRs using the default Martini 3.0.0 force field result in too compact IDR conformations and an underestimation of the radius of gyration^10,13,36,37^. Therefore, the force field should be modified to account for the specific properties of IDRs. Here, we present two options: modifying the BB-water interaction strength and the dedicated force field Martini3-IDP.

### 4.1 Adding a Water Bias

One way to improve the performance of Martini 3 when simulating IDRs is to modify the strength of the protein-water interactions for the IDR only. This technique utilizes the BB virtual sites, which are also used in the GōMartini model to implement the structural bias, to modify the LJ potential depth between the BB and water beads. Since these virtual sites overlap with the BB bead, the non-bonded interactions between BB and water beads can be effectively adjusted by introducing an additional potential between the virtual site and water beads^13^.

In this approach to set up the multi-domain protein model, we use the full length CK1δ atomistic structure as Martinize2 input. As the IDR structure is typically not present in experimental structures, AlphaFold or SWISS-MODEL can be used to generate the full length atomistic protein structure (*CK1d_IDR*.*pdb*)^23,24^. In Martinize2, the water bias can be added via the *-water-bias* flag. The residues of the IDR are defined via -*id-regions 293:415*, and the strength of the water-bias is set to a value of 0.5 kJ/mol here by *-water-bias-eps idr:0*.*5*^13^. Additionally, we need to manually specify the secondary structure of our protein with the flag *-ss* using the string returned by DSSP in Section 2.3.1. To avoid creating unwanted secondary structures in the IDR of the protein, which are sometimes wrongfully present in the atomistic reference structure, replace all characters of the IDR with ‘C’ for coil. The water bias can be combined with the GōMartini model for the structured region, although any of the protein structure models discussed previously are also compatible. Here we add an EN for the structured region with *-eunit 1:292*.

~~~
martinize2 -f CK1d_IDR.pdb -x CK1d_cg.pdb -o CK1d_only.top -ff martini3001
-p backbone -ss CCCEETTTEEEEEEEEEETTEEEEEEEETTTTEEEEEEEEETTSSSCCHHHHHHHHHHHTTCTTCCE
EEEEEETTEEEEEEECCCCBHHHHHHHTTTCCCHHHHHHHHHHHHHHHHHHHHTTEECSCCCGGGEEECCGGGTTCEEECCG
GGCEECBCTTTCCBCCCCCSCCCCSCTTTCCHHHHTTCCCCHHHHHHHHHHHHHHHHHSSCTTSSCCCSSHHHHHHHHHHHH
HTSCHHHHTTTSCHHHHHHHHHHHHCCTTCCCCHHHHHHHHHHHHHHTTCCCSCCCCCCCCCCCCCCCCCCCCCCCCCCCCC
CCCCCCCCCCCCCCCCCCCCCCCCCCCCCCCCCCCCCCCCCCCCCCCCCCCCCCCCCCCCCCCCCCCCCCCCCCCCCCCCCC
CCCCCCCCCCCCCCCCCCC -elastic -el 0 -eu 0.85 -eunit 1:292 -water-bias -water-bias-
eps idr:0.5 -id-regions 293:415
~~~

In addition to the CG protein structure and topology, Martinize2 returns two files: virtual_sites_atomtypes.itp, and virtual_sites_nonbond_params.itp, which need to be added to the Martini 3 topology file, as described in Section 2.3.3.

### 4.2 Martini3-IDP

Another way to improve the performance of IDRs in Martini 3 is offered by the recently developed Martini 3 disordered protein force field (Martini3-IDP)^38^, which is well integrated into the current Martini 3 framework. All force field files and Python scripts in this tutorial Section are available at https://github.com/Martini-Force-Field-Initiative/Martini3-IDP-parameters.

Currently, two methods to generate the Martini3-IDP model are available.

#### 4.2.1 Setup Using Martinize2 and Polyply

In this approach, the Martini model of a folded domain and the IDR are first generated separately with Martinize2 and Polyply. In a second step, both models are merged.

As in Section 2.3.2, the Martini model of the CK1δ protein folded domain with EN bias is generated by Martinize2 as following. The EN bias is used in this tutorial Section as example, but GōMartini and OLIVES could also be transferred easily referring to the tutorial in Section 2.3.

~~~
martinize2 -f CK1d.pdb -x CK1d_cg.pdb -o CK1d_only.top -ff martini3001 -p backbone
-dssp -elastic -el 0 -eu 0.85
~~~

Polyply is used to generate the topology and coordinates files of the IDR and can be obtained from GitHub (https://github.com/marrink-lab/polyply_1.0)^39^. *polyply gen_params* is used to generate the topology with protein sequence fasta input by *-seqf* (or protein sequence string by *-seq*). The *-f Martini3-IDP_Polyply*.*ff* flag provides the force field. *polyply gen_coords* is used to generate random IDR coordinates file by a random walk. As input, only the IDR topology is required (*-p* flag). The flag *-box* defines the edge length of the simulation box, *-o* defines the output file name containing the coordinates.

~~~
polyply gen_params -name IDR -f Martini3-IDP_Polyply.ff -seqf IDR.fasta -o IDR.itp
polyply gen_coords -p topol.top -name IDR -box 30 30 30 -o IDR.gro
~~~

Note that the topology provided with the *-p* flag is not the .itp file of the IDR, but the GROMACS .top file including all necessary force field files involved in *IDR*.*itp*, similar to this:

~~~
#include “./martini_v3.0.0.itp”
#include “./martini_v3.0.0_solvents_v1.itp”
#include “./martini_v3.0.0_ions_v1.itp”
#include “IDR.itp”
[molecules]
IDR 1
~~~

Finally, a Python script *topology_merging_general*.*py* is used to merge all domains’ .itp files and define the parameters of the crosslinking region. You must change the .itp file name and molecule type to “mol0 mol1 …” according to their order in the protein. And because sometimes dihedral parameters in Martini topology are defined in two individual [dihedrals] derivatives, for consistency, we need to merge [dihedrals] derivatives in the .itp files with two [dihedrals] derivatives by definition in *topology_merging_general*.*py*.

Then based on the domain number in the protein, we need to adjust some definitions by order in *topology_merging_general*.*py* (For example, two domains here in CK1δ protein “mol0-mol1” are merged).

~~~
mol0_BB, mol0_SC, mol1_BB, mol1_SC = entry_edit(mols, ‘mol0’, ‘mol1’)
interaction_adding(ff, molname, mol0_BB, mol0_SC, mol1_BB, mol1_SC)
~~~

Because the protein CK1δ contains only one folded domain and one IDR, ‘mol0-mol1’ is specified with the *-i* flag. The default name of the output .itp file is *mol*.*itp*.

~~~
python3 topology_merging_general.py -i mol0-mol1
~~~

Note that the terminal bead type in each domain fragment (i.e. the crosslinked residues in the final multi-domain protein) must be checked and adjusted manually to the default Martini 3 backbone bead for that particular residue in the output *mol*.*itp* (see Figure 1).

~~~
714 P2 292 MET BB 714 0.0
⁝
716 P2 293 LEU BB 716 0.0
~~~

To obtain the coordinate file of the complete multi-domain protein, insert separate domains structures into one box, then after a short simulation, the broken domain fragments should be connected due to the bonded linking between fragments in *mol*.*itp*.

#### 4.2.2 Setup Using Martinize2 only

A Martinize2 format Martini3-IDP force field file is also provided, which deals with the differences between the standard Martini 3 protein model and Martini3-IDP model. For example, *-scfix* is applied in the Martini 3 protein model to define the sidechain orientation relying on atomistic reference structure, whereas general bonded parameters are used to define sidechain orientation in Martini3-IDP independent on the atomistic reference structure.

Similar to Section 4.1, the EN bias is generated only for the folded domain using the *-eunit* flag. To define the IDR easily, a new secondary structure ‘D’ is provided in the -ss flag. Note that ‘D’ is not a pre-set secondary structure, so ‘D’ secondary structure should be manually added to ‘vermouth/dssp/dssp.py’ ss_cg list.

~~~
martinize2 -f CK1d_IDR.pdb -x CK1d_cg.pdb -o CK1d_only.top -ff martini3001-IDP
-p backbone -ss CCCEETTTEEEEEEEEECSSSEEEEEEETTTTEEEEEEEEETTCSSCCHHHHHHHHHHHTTSTTCC
CEEEEEEETTEEEEEEECCCCBHHHHHHHTTTCCCHHHHHHHHHHHHHHHHHHHHTTEECCCCCGGGEEECCGGGTTCEEEC
CCTTCEECBCTTTCCBCCCCBSTTCCSCTTTCCHHHHTTBCCCHHHHHHHHHHHHHHHHHSSCTTSSCCCSSGGGHHHHHHH
HHHHSCHHHHTTTSCHHHHHHHHHHHHSCSSCCCCHHHHHHHHHHHHHHTTCCCSCCCGGGCDDDDDDDDDDDDDDDDDDDD
DDDDDDDDDDDDDDDDDDDDDDDDDDDDDDDDDDDDDDDDDDDDDDDDDDDDDDDDDDDDDDDDDDDDDDDDDDDDDDDDDD
DDDDDDDDDDDDDDDDDDDDD -elastic -ef 700.0 -el 0 -eu 0.85 -eunit 1:292
~~~

In the near future, the Martini3-IDP model will be fully implemented in Martinize2, simplifying the generation of multi-domain protein models with Martini 3.

### 4.3 Simulation of the Multi-Domain CK1δ Model

As described in Section 2.4, the system can be solvated using *insane* with the following command.

~~~
insane -f CK1d_cg.pdb -o CG.gro -p system.top -pbc cubic -box 35,35,35 -salt 0.15
-sol W -d 0
~~~

When simulating a protein with an IDR, it is especially important to use an appropriately sized box, to prevent the extended IDR from interacting with its mirror image. We therefore use an edge length of 35 nm here. The simulation of the system follows the protocol outlined in Section 2.4.

The conformational ensemble sampled during the simulations is shown in Figure 10.

**Figure 10:**
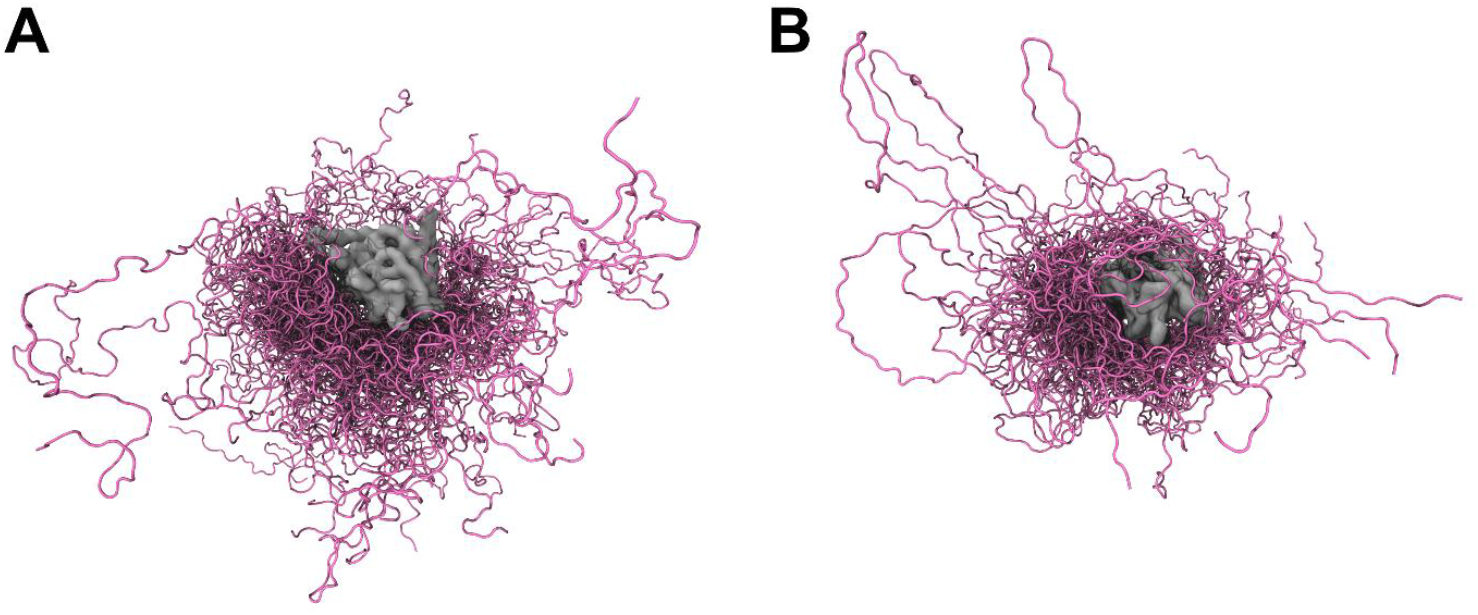
Simulation of CK1δ including the IDR using the modified water bias (A) and the Martini3-IDP model (B). Displayed are 100 frames of a 5 μs simulation, with the structured region shown in gray and the IDR in pink.

## 5 Notes and Limitations

Before summarizing, we would like to provide some additional notes to the reader pointing to advanced aspects of Martini 3 protein models. Moreover, we will discuss its most important limitations.

### Note 1 - Further optimization of protein structures

In Section 3.1.1, it was demonstrated how the biasing layer of Martini proteins could be adjusted by changing global settings such as cut-off or bias strength. However, spending more time on optimization, you could tune these aspects also on a local basis, i.e., by removing or adjusting only part of the biasing network. For instance, to improve regions that are too rigid or too flexible in comparison to all-atom reference simulations (as measured by, e.g., RMSF analysis), or to improve the shape fluctuations of a binding pocket or entrance pathway. Although in principle this can often be done manually, the MAD server offers a graphical interface for adding or removing bonds (https://mad.ens-lyon.fr/)^40^.

### Note 2 – Multiple reference structures in one local minimum

In Section 2, a single reference structure was used to generate the protein model. Potential artifacts and rare conformations of this reference, such as specific conformations of flexible loops or side chain orientations influenced by the crystal packing, are thus incorporated in the model. One way to improve this is to use multiple reference structures. Based on the contact frequency within the set of reference structures, selected contacts can be removed from the structure bias model if they do not occur in the majority of the reference structures. These can be either from experiment, for instance NMR structures, or from atomistic reference simulations. The latter has been shown to improve in particular the flexibility of loops for an exemplary test set of six proteins with the GōMartini 3 protein model^13^. The OLIVES approach currently allows to use multiple references for the generation of the contact map^17^.

### Note 3 – Reference structures from two conformational states

The multiple-basin GōMartini approach combines two GōMartini 3 models for two distinct states of the protein using an exponential mixing scheme^41^. This extension offers the possibility to investigate conformational transitions of proteins and to identify intermediate conformational states. Note that the program package OpenMM^42^ is required for the multiple-basin GōMartini approach. A simpler way of modelling conformational transitions is to switch between two GōMartini models^43^. Reference structures from two conformational states can also be used to generate an OLIVES model^17^.

### Note 4 – OliGōmers

An extension to the GōMartini 3 model named OliGōmers offers a computationally efficient way to model protein oligomers using the Gō-like model as structure bias model^44^. Multimeric proteins require the definition of separate .itp files for each monomer with the GōMartini 3 model. This is necessary because the virtual sites which take care of the Gō-like interactions do not distinguish between the intra- and intermolecular interactions. Thus, large oligomers such as fibrils are challenging to describe. This issue is solved via a multi-layer virtual site scheme in the OliGōmers implementation.

### Note 5 – *In silico* single molecule force spectroscopy

The use of Gō-like models as structure bias models for Martini proteins opened the way to efficient single molecule force spectroscopy calculations^13,18,45,46^. The key feature of these Martini protein models was the ability of protein unfolding due to breakable Gō-like interactions. To faithfully reproduce experimental force-extension data of protein complexes, it is often required to add Gō-like interactions at the protein-protein interfaces^13,47^.

### Note 6 - Visualization of protein structures

Most visualization packages, including the popular VMD^48^, cannot draw correct bonds based on Martini structure files only, and require loading of topology files as well. However, many Martini models (including proteins) now make extensive use of interaction types like virtual sites, requiring a dedicated script to allow VMD to correctly depict the CG topology. MartiniGlass is such a script, freely available on GitHub (https://github.com/Martini-Force-Field-Initiative/MartiniGlass)^49^. The program has a particular focus on being able to visualize protein secondary/tertiary structure networks, although MartiniGlass can in fact be used to reconstruct bonded networks of any Martini molecule.

### Limitation 1 – Changes in secondary structure

Currently, the Martini 3 protein model is not able to capture folding processes and cannot fully capture changes in secondary structure – at least using the standard protocols to build the CG protein models. One of potential reasons for this limitation is the lack of directionality, which is important for certain interactions such as hydrogen bonds and T-shaped interactions of aromatic units^6^. In addition, some of the bonded terms of the current Martini 3 protein model depend on the secondary structure, if secondary structure information from DSSP is used for the model. One promising development towards secondary structure changes with respect to Martini 2 proteins is, however, that the particle type of the backbone is not secondary structure-dependent anymore^10,13^.

### Limitation 2 – Protonation states and Martini sour

In standard molecular dynamics protocols, it is common to assume a fixed protonation state of titratable groups such as the side chains of arginine, glutamic acid, and histidine. However, this is not always realistic, for instance if the pH is close to the pK_a_ value of the titratable group or if the environment modulates its pK_a_. Constant pH approaches or the titratable Martini model offer ways to introduce changes of protonation states^50–52^. In titratable Martini, a dedicated class of beads, so-called titratable beads, can bind or release a proton particle mimicking H^+^. The titratable Martini model and constant pH approaches pave the way to study the impact of pH on protein conformations^53^.

### Limitation 3 – Effective time scale

Inherent to coarse-graining is the loss of degrees of freedom. This smoothens the potential energy surface and results in an effective time scale for CG simulations, which complicates direct comparison to experimental and atomistic simulation data. A challenge is that the scaling factor between real time and the effective CG time is system dependent. Here, we estimated a scaling factor of 4 to compare atomistic and CG data^17,34^. However, please be aware that this is just an estimate and no universal scaling factor for Martini 3.

### Limitation 4 – Hydrophilicity of Martini 3 proteins

In the Martini 3 protein model, the backbone is mapped to one bead type independent of the secondary structure. This is a step towards a protein model which is able to change the secondary structure. However, this entails that the partitioning of the backbone cannot be used anymore to compensate possible differences in hydrophilicity of secondary structure elements. There have been reports of too hydrophilic single-pass transmembrane helices and β-sheet dimerizing peptides^54,55^, and too hydrophobic intrinsically disordered proteins^36,56,57^. Adapting the backbone-water interactions (see Section 4.1) and refining the dihedral potentials in the Martini-IDP force field, respectively, offer a way to improve the behaviour of Martini 3 proteins. Note that alternative approaches such as rescaling of the full protein-protein or protein-water interactions have to be taken with care ^36,54,56,57^, because they also modify the well-balanced side chain partitioning^13^.

### Limitation 5 – Non-canonical amino acids and post-translational modifications

While simple post-translational modifications such as disulfide bridges, phosphorylation, and capped terminals can be straightforwardly included in Martini 3 protein models via Martinize2^15^, more complex modifications remain a challenge. Lipidation is available in Martini 3^58^, as it was in Martini 2^59^, and PEGylation can be incorporated using Polyply^20,39^. However, glycosylation is still under development, alongside with the development of carbohydrate models^60,61^. Furthermore, differences in chirality, such as modelling D-amino acids, are not yet supported, as current protein models only consider L-amino acids. The same applies to other chemical, non-biological modifications often used in the design of biomimetic peptides. These limitations restrict the ability to study non-canonical and heavily modified proteins without extensive manual parameterization or specialized tools

## 6 Summary and Outlook

In this chapter, we introduced the reader to setting up and characterizing a Martini 3 protein model with different structure bias models, namely Elastic Network^3,15,16^, GōMartini^13^, and OLIVES^17^. In addition, we introduced two options to model IDRs and to model multi-domain proteins with structured and intrinsically disordered regions^13,62^. Finally, we discussed some advanced aspects of modelling CG proteins as well as limitations of the current Martini 3 protein model.

The Martini 3 protein model represents a significant advancement in coarse-grained molecular dynamics, offering improved flexibility and accuracy compared to its predecessors. With its ability to model a wide range of biomolecular systems, including structured and intrinsically disordered proteins, Martini 3 opens the door to studying complex biological phenomena on time and length scales inaccessible to atomistic simulations, extending its potential toward simulations of entire organelles and even cells^63,64^. However, continued efforts are required to address existing limitations, such as modelling structural changes ranging from local dynamics to global transitions, including the accurate representation of multiple folded states of the same protein. Additionally, further developments are needed to incorporate non-canonical amino acids and fully integrate diverse post-translational modifications. Advances in tools like Martinize2^15^ and Polyply^39^, together with ongoing developments of improved models like Martini3-IDP^62^, promise to expand the applicability of Martini 3 to even more diverse systems, including biomimetic designs and systems influenced by dynamic environmental conditions like pH. As these developments unfold, Martini 3 is poised to become an even more powerful tool for researchers across biophysics, drug development, and biomaterials design, bridging the gap between detailed atomistic insights and large-scale biological phenomena.

## 7 Acknowledgements

T.D. and S.T. thank the Alfons und Gertrud Kassel Foundation, the Dr. Rolf M. Schwiete Foundation, the H. & E. Kleber Foundation, the SCALE cluster of excellence initiative, and the Center for Multiscale Modelling in Life Sciences (CMMS) sponsored by the Hessian Ministry of Science and Art for funding and the Center for Scientific Computing at Goethe University Frankfurt for access to Goethe-HLR. L.W. was supported by the China Scholarship Council. L.B.A. and P.C.T.S. acknowledge the support provided by the CNRS. They also acknowledge the support of the Centre Blaise Pascal’s IT test platform at ENS de Lyon (Lyon, France) for the computer facilities. The platform operates the SIDUS solution developed by Emmanuel Quemener^65^.

